# Light-Dependent Ethanol Assimilation By Model Red Microaga *Galdieria sulphuraria*

**DOI:** 10.1101/2024.11.22.624789

**Authors:** Igor Stadnichuk, Yulia Bolychevtseva

## Abstract

The effect of ethanol on the red thermoacidophilic microalga *Galdieria sulphuraria*, capable of hemo- and photoheterotrophic growth, was studied. It was found that *G. sulphuraria* failed to grow and assimilate ethanol when kept in the dark, in contrast to a wide range of organic compounds known to this alga. Light conditions, on the other hand, activated ethanol uptake from the growth medium and intensify microalgal growth. The known stress effect of ethyl alcohol can be seen in the short-term increase in cellular respiration in the dark and its sustained increase throughout the growth period of *G. sulphuraria* in the light. Ethanol-induced increase in cellular respiration precedes light activation of photosynthesis. The growth enhancement of ethanol-supplemented culture of *G. sulphuraria* compared to photoautotrophic culture is most likely due to the complete oxidation of ethanol during respiration to carbon dioxide and its use by chloroplasts as an additional intracellular substrate for photosynthesis. The absence of ethanol consumption in the dark and its metabolic uptake in the light indicate photoactivation of alcohol dehydrogenase and acetaldehyde dehydrogenase, two key sequential enzymes in the ethanol oxidation pathway.

## Introduction

Species of the genus *Galdieria* belong to the phylum Cyanidiales, a group of red algae that has been isolated early in evolution (Yoon et al., 2006). Most of its members inhabit geothermal sulphurous springs containing heavy metal ions at pH 0-4 and temperatures up to 56°C (Seckbach, 1981). These microalgae combine eukaryotic cell structure with ecological and biochemical characteristics of extremophilic prokaryotic species. Due to their exceptional properties and the minimal genome size of 12-18 Mb (Muravenko et al., 2001; Miyagashima et al., 2021; Stadnichuk and Tropin, 2022). Cyanidiales as representatives of archaeoplastids have become one of the most important objects of research in cell physiology, biochemistry, molecular biology, phylogenomics and evolutionary biology. Polyextremophilia and many metabolic traits have been inherited by Cyanidiales from archaea and bacteria, demonstrating the possibility of horizontal gene transfer from prokaryotes to eukaryotes (Yoon et al., 2006; Stadnichuk and Tropin, 2022).

In addition to photoautotrophy, the genus *Galdieria* is characterised by the ability to grow both dark (chemoheterotrophic) and light (mixotrophic) heterotrophic, known to occur in several groups of unicellular algae (Selosse et al., 2017; Pribyll and Cepak, 2019). Heterotrophy serves as an adaptive strategy that allows the use of dissolved organic matter when it becomes the most favourable energy source in the microalgal habitat (Selosse et al., 2017). Thus, the possibility of photo- and chemoheterotrophic growth is a successful evolutionary strategy of microalgae that increases their physiological and biochemical activity and adaptive capabilities (Selosse et al., 2017). Twenty-seven digestible carbohydrates and polyols and, taking into account a number of saccharophosphates, amino acids and Krebs cycle intermediates, up to 50 exogenous organic substrates have been identified that can be used by species of the genus *Galdieria*, which is a kind of record for microalgae (Gross and Schnarenberger, 1995; Oesterholt et al.; Schmidt et al., 2005; Yoon et al., 2006; Pribyll et al., 2019;). The model species *Galdieria sulphuraria* owes its ‘omnivorousness’ and metabolic plasticity to the presence of appropriate transfer proteins and enzymes for the assimilation of mono- and disaccharides and polyatomic alcohols from the external environment (Barbier et al., 2005; Ternes, 2015; Fujiwara et al., 2019). However, the relationship between respiratory and photosynthetic processes in photoheterotrophy still leaves many uncertainties that require further studies of *Galdieria* genus and other algal mixotrophs (Gross and Schnarenberger, 1995; Tischendorf et al., 2007; Selosse et al., 2017).

Extremophyles are of increasing interest due to their unique metabolic abilities and great biotechnological potential. Bioremediation of domestic and industrial wastewaters containing organic matter and having elevated temperature serves as the most promising direction for the use of microalgae of the genus Galdieria in biotechnology. Coastal algal mats of hot springs containing *G. sulphuraria* indicate the possibility of immobilisation of *Galdieria* strains in wastewater treatment plants (Lang et al., 2020). Therefore, the use of microalgae of the genus *Galdieria* for neutralisation of food organic waste looks very promising (Selvaratnam et al., 2014; Cizkova et al., 2019).

The list of exogenous substrates for *Galdieria* is extensive, but not exhaustive. Ethanol is not mentioned among these organic products, although ethanol may even serve as an additional energy source in the process of neutralising ethanol stress. Ethanol is a natural product of carbohydrate fermentation in yeasts of the genus *Saccharomyces* and bacteria *Zimomonas*, and is common in fermenting berries and fruits. It was important to investigate the effect of ethanol on a model species, *G. sulphuraria*, in relation to the use of food waste, of which fruit and berries can be a significant proportion. Ethanol is toxic and, because of its low degree of dissociation, is easily soluble in lipids and water. Alcohol readily enters cells and tissues and leads to disoder of biomembranes, changes in the conformation of proteins and a cascade of metabolic disturbances, including the appearance of reactive oxygen species (Uspensky, 1984; Bullock, 1990; Averina et al., 2020). Ethanol neutralisation is achieved in all organisms by its oxidation to harmless products - carbon dioxide and water. The universal and evolutionary oldest pathway, starting with archaea and bacteria, is ethanol metabolism by alcohol dehydrogenases (ADH) (Fig. 1) (Kornmann et al., 2003). In humans, in addition to the main ADH pathway, a small fraction of ethanol is converted by peroxidation with aldolase and the cytochrome P450 pathway (Uspensly, 1984; Bullock, 1990). In green microalgae and dinoflagellate species that have been studied in relation to ethanol, as well as in cyanobacteria as chloroplast precursors, ethanol conversion occurs via ADH (de Swaaf et al., 2003; Vidal et al., 2009; Wen et al., 2015 et al; Jiang., 2017; Lang et al., 2020) but in *Euglena gracilis*, in addition to ADH, a microsomal system of alcohol oxidation via the alcohol oxidase pathway similar to that in humans has been found (Palma-Gutierrez et al., 2008).

**Fig. 1.**
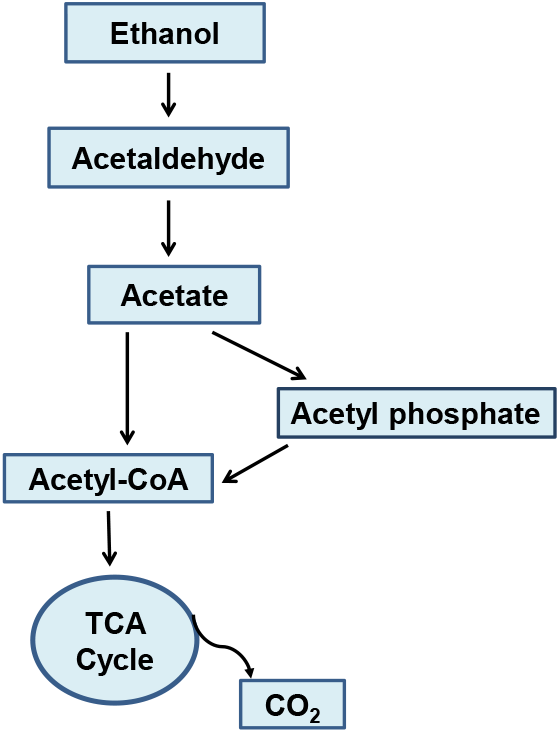
Metabolic pathway for oxidative assimilation of ethanol into the TCA cycle via Acetyl-CoA

On the other hand, ADH is known to be a bi-functional protein that, along with ethanol, can neutralise other cellular stress factors such as reactive oxygen species. This may explain the early evolutionary appearance of this enzyme. It is therefore not surprising that a system of active ethanol metabolisation by ADH may have already evolved in unicellular organisms.

The aim of this work was to find out the effect of ethanol on the growth of the microalga *G. sulphuraria*. It was shown that *G. sulphuraria* is able to metabolise ethanol in the light, which leads to increased photosynthetic activity and intensified growth of the culture, whereas in the dark, in the absence of photosynthesis, ethanol is not metabolised.

## Matherials and Methods

### Strain and culture conditions

The axenic culture of the thermoacidophilic red microalga *Galdieria sulphuraria* (Galdieri) Merola, strain IPPAS P–513, from the IPPAS microalgae and cyanobacteria collection (RAS) was used. Allen’s growth medium was used with the pH reduced to 2.5 by micro-addition of H_2_SO_4_ [8]. Algae were grown at 38°C in Allen’s medium, with the pH adjusted to 2.5 by micro-addition of H_2_SO_4_ (Gross and Schnarenberger, 1995). The photoautotrophic culture and the culture containing 0.1% ethanol were grown at 38°C under 24 h constant white light of 40 μM photons-m^−2^ s^−1^ (SL 20/32–735 lamps, China). 100ml conical flasks were used, each containing 50ml culture medium (4cm liquid layer). Cell suspensions were manually agitated daily for 5 min without bubbling or shaking to reduce ethanol evaporation (Dubock and von Stockar, 1998).

### Cell growth curves and absorption spectra

Cell suspension samples were taken five times: on the day of inoculation and then on days 2, 5, 9 and 11 of growth in three biological replicates. Further measurements were stopped due to increased evaporation of ethanol from the culture medium at 38 °C. Selected aliquots of 1.5 ml were used to measure light scattering and absorption spectra. The absorption spectra of the samples were obtained in 1.0 cm optical cuvettes on a Beckman DU-650 spectrophotometer (“Beckman Coulter Inc.”, USA) in the spectral range 400-750 nm. The level of scattered light due to minimal absorption of chlorophyll and other photosynthetic pigments in the near-infrared region at 750 nm was proportional to the dry mass of cells accumulated in suspension, providing a simple and convenient way to construct growth curves (Ketsuda et al., 2006; Schwerna et al., 2016). To improve the accuracy of calculations, the proposed formula (Ketsuda et al., 2006) was used, taking into account the absorption intensity of chlorophyll at 678 nm in addition to the spectral data at 750 nm. In order to obtain spectra with a correct representation of the absorption properties of the pigment apparatus, the contribution of scattered light in the entire spectral region of registration was subtracted from the absorption spectra of the samples using the mathematical software ‘Origin6.1’.

### Ethanol losses

Cells were sedimented by centrifugation for 7 min at 7000 g in an Eppendorf centrifuge (MiniSpin, Germany). The ethanol content of 1.5 ml supernatant samples was determined chromatographically at the same times as used to assess culture growth. A Crystal 5000.2 gas chromatograph (‘Chromatek’, Russia) was used. It was equipped with a ZB-WAXplus 30 m x 0.25 mm x 0.35 μm capillary column (‘Phenomenex’, USA) and a PID detector. The carrier gas was high purity nitrogen (EP grade). Separation was performed in temperature gradient. Cell-free control samples with added ethanol were used to account for the increased evaporation of alcohol from the culture medium (Dubock and von Stockar, 1998) at 38 °C throughout the experiment. (Rectified Ethyl Alcohol, ‘Lux’, 96.3%, GOST 5962-2013, LLC ‘Donskoy’, RF). All other chemicals used wer of guaranted grade from commercial sources.

### Cellular respiration and photosynthesis

Oxygen uptake in the dark and oxygen release in the light were recorded directly in a suspension of *G. sulphuraria* cells on a potentiometer equipped with a Clark electrode (Oxygraph, “Hansatech”, Germany) as described in (Voloshina et al., 2016; Schwerna et al., 2019) in a thermostated 1 mL cuvette at 38 °C. The natural electron donor (H2O) and acceptor (CO_2_) were used for the determination of photosynthetic activity. Cell suspensions were preconcentrated to equal optical scattering level values of OD_750_ by gentle filtration (Nalgene filters, pore size 0.45 μm). The change of oxygen concentration in the medium was expressed in nmol O_2_/(OD_750_) ml min. The source of white light was an OI-24 illuminator (LOMO, Russia) with an illumination intensity of 800 μM photons s^−1^ m^−2^. Immediately prior to measurements, the harvested cells were incubated for 30 min in the dark to adapt to the dark conditions and for 20 min in the white light of the SL 20/32–735 lamp used for the culture to pre-adapt to the light.

### Statistical Analysis

Data were analyzed using one-way ANOVA tests, with *p* < 0.05 denoting significance.

## Results

### Growth curves

The ability of microalgae to assimilate ethanol is assessed by changes in biomass growth during cultivation. Due to the toxicity of ethanol, unicellular photosynthetics stop growing or even die when the ethanol content exceeds 4%. The content of added ethanol in growth experiments with microalgae can start as low as 0.1 % and usually does not exceed 0.5 - 1.0 % (Bozhkov et al., 2014; Jiang et al., 2017). Optimal concentrations are species-specific. For example, the biomass of the green alga *Dunaliella viridis* decreases in the presence of 0.5% ethanol in the medium compared to its fraction equal to 0.3% (Schwerna et al., 2016), but in the dinoflagellate *Crypthecodinium cohnii* the concentration of 0.5% is optimal (de Swaaf et al. 2003). Only in *E. gracilis*, ethanol remains an active growth stimulator up to a concentration of 1.5% (Rodriguez-Zaval et al., 2006). The addition of ethanol to the culture of the green alga *Chlamydomonas reinhardtii* caused an induction or repression of the biosynthesis of up to 40% of different proteins depending on the alcohol concentration (Jiang et al., 2017) In this regard, there is a concept of ‘mild stress’ for small amounts of incoming ethanol (Bozhkov et al., 2014). This allows an ethanol metabolising cell to gain energetic benefits from low concentrations while minimising the toxic effects of ethanol. For these reasons, ethanol as a substrate occupies a special place in the list of organic products consumed.

In the present work, the possibility of *G. sulphuraria* growth in the presence of ethanol in light and dark was studied. A minimum concentration of 0.1% (21.7 mM, w/v) was used to reduce ethanol stress in all experiments performed. It was preliminarily observed that, in the light, increasing the ethanol concentration to 0.5% gave virtually no increase in growth (data not shown) compared to a concentration of 0.1%. Under dark conditions, the same concentration was used. Fig. 2 demonstrates the growth curves of *G. sulphuraria* culture under photoautotrophic conditions without ethanol supplementation and in its presence in light and dark.

**Fig. 2.**
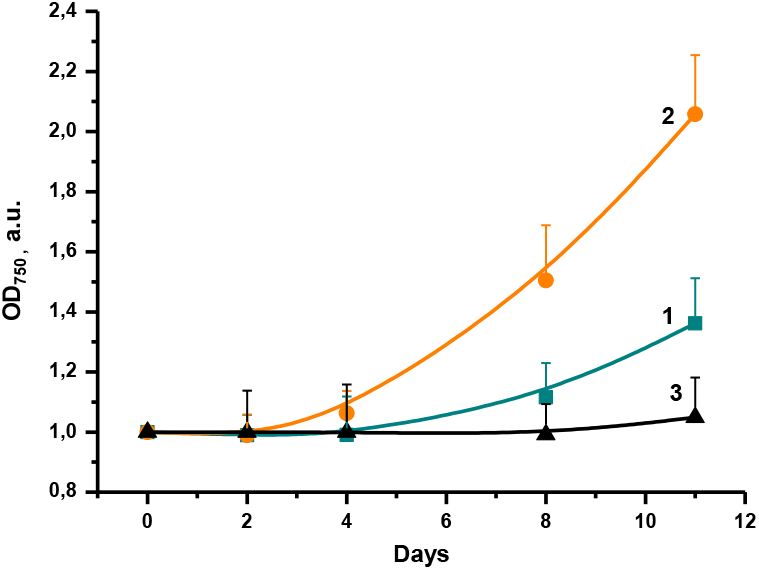
Growth curves of *G. sulphuraria* cell culture placed in light or dark in the presence or absence of ethanol. 1 Slow growing under photoautotophic (light only) conditions. 2 rowth enhancement with addition of 0.1% ethanol under light conditions. 3 Failure to increase growth by supplementing the culture medium with 0.1% ethanol in the dark. Curve 3 corresponds exactly to the survival curve not shown for a darkened autotrophic culture.

The peculiarity of the photoautotrophic culture of G. sulphuraria is its extremely slow growth, as a considerable part of the energy used by the cells is spent on the pumping out of protons penetrating from the medium and the maintenance of the neutral intracellular pH at its external value of 2. The buffer equilibrium 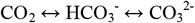 at pH <4 is shifted completely in favour of CO_2_, while the solubility of carbon dioxide is significantly reduced at elevated temperature, which also results in low photosynthetic activity and slow photoautotrophic growth (Stadnichuk and Tropin, 2022). At the end of the experiment on day 11, the photoautotrophic culture grows only about one and a half times, but in the presence of ethanol it more than doubles. No statistically significant changes in growth were also observed in the dark in the presence of ethanol, as opposed to light (Figure 2). This is an important difference from known non-toxic organic substrates, because in the presence of glucose or other hexoses and glycerol, not only photoheterotrophic but also dark heterotrophic growth of *G. sulphuraria* is observed, which is far superior to photoautotrophy (Gross and Schnarenberger, 1995; Oesterhelt et al., 1999). Microscopic control showed that the cells in both dark cultures remained viable, i.e. in a state of survival, throughout the duration of the experiment.

### Ethanol Loss

Ethyl alcohol is a readily evaporating liquid with at least 20% loss (Duboc and von Stockar, 1998) during stationary cultivation at room temperature. Under our conditions, at 38 °C, the evaporation is even more pronounced, as shown by the chromatographic determination of the alcohol loss from the culture medium. It amounts to 65% on day 11, towards the end of the incubation (Fig. 3).

**Fig. 3.**
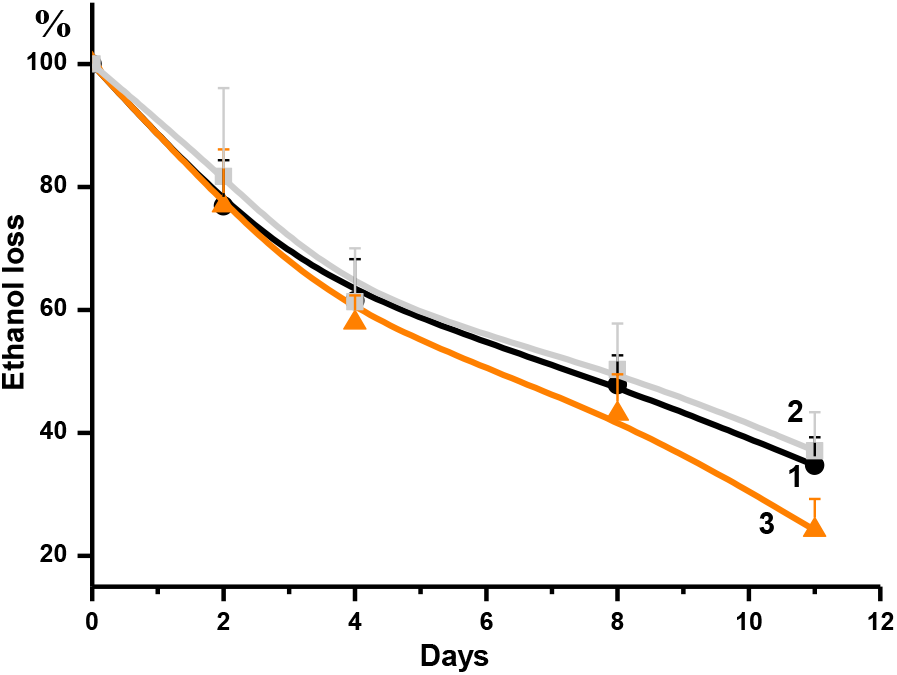
Loss of ethanol from the culture medium during *G. sulphuraria* incubation period. 1 Evaporation of ethanol in the absence of *G. sulphuraria* cells. 2 No difference from curve 1 when *G. sulphuraria* cells were in the dark with 0.1% ethanol. 3 Additional ethanol lost due to *G. sulphuraria* using it in the light.

Curve 2, obtained for the ethanol-containing cell culture grown in the dark, is almost identical to curve 1 within the accuracy of the experiment at all times of measurement, confirming the non-use of alcohol by *G. sulphuraria* cells under dark conditions (Fig. 2). Additional loss of alcohol is revealed only in the light. By the end of the experiment, ethanol level decreases by another 26% compared to the level maintained by simple evaporation. Thus, the non-usage of ethanol by *G. sulphuraria* cells in the dark and the consumption of ethyl alcohol from the culture medium in the light correlate with the data on additional light-induced cell growth (Fig. 2).

### Respiration and photosynthesis of *G. sulphuraria* under light conditions

The functional activity of *G. sulphuraria* cells grown in the light was assessed by measuring the intensity of photosynthesis compared to cellular respiration. In the photoautotrophic growth regime, cell respiration remains low and its slight increase during the experiment is at the limit of measurement accuracy and can be attributed to an increase in the frequency of cell division (Fig. 4a, columns A). In the presence of ethanol, there was a significant increase in oxygen consumption as early as day 2 (B* > B) and a gradual increase thereafter. It only decreased slightly on the last day of the experiment, probably due to a marked decrease in the ethanol content of the culture medium (see Fig. 3). Thus, on all the days of the experiment, starting from the second day, the level of respiration during the addition of ethanol was several times higher than in the photoautotrophic culture (B* > A).

**Fig.4.**
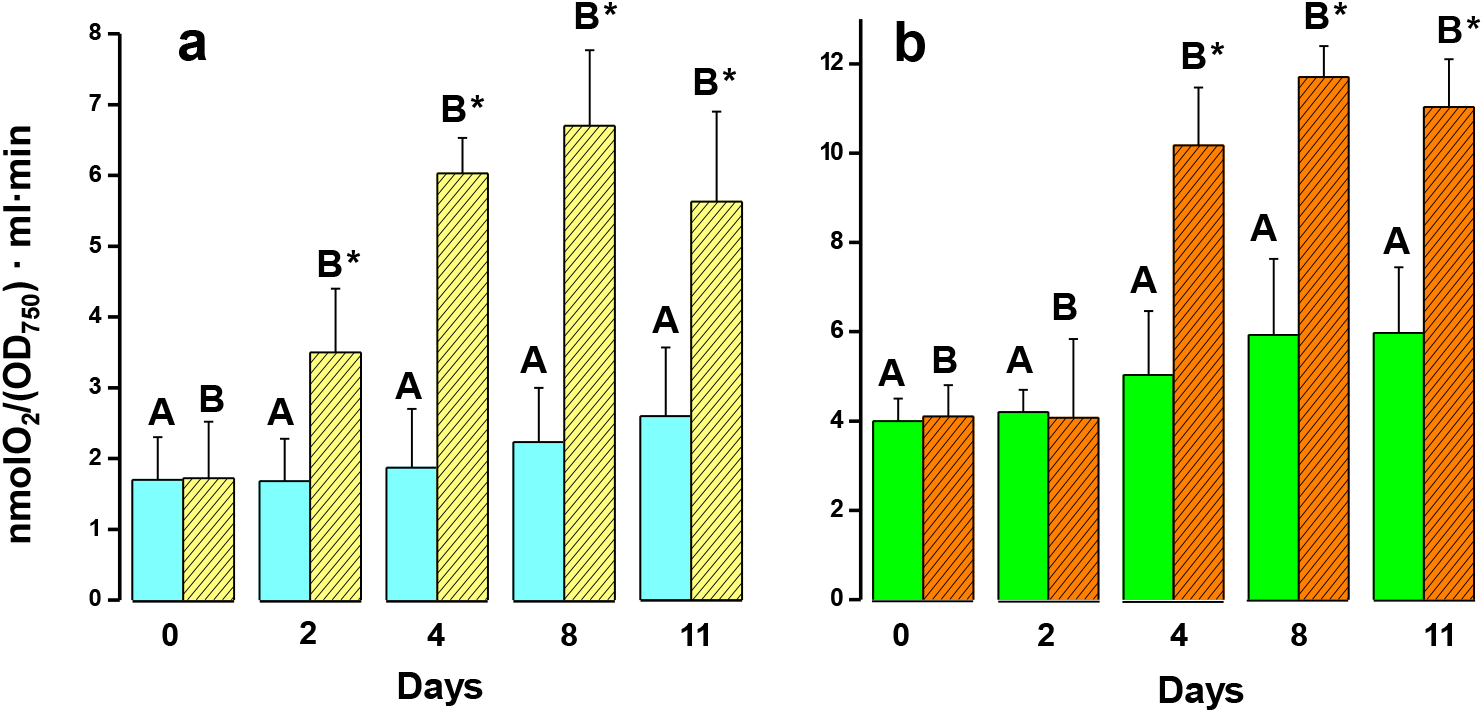
Oxygen uptake during dark respiration (a) and oxygen release during photosynthesis corrected for respiratory uptake of O_2_ (b) for *G. sulphuraria* culture. Columns A correspond to photoautotrophic growth and columns B to the addition of 0.1% ethanol. Statistically significant differences are marked with *.

Determination of photosynthetic oxygen evolution in the photoautotrophic culture showed that, judging from the mean values, there has been a slight increase, similar to the increase in respiration. Compared to photoautotrophic growth, the enhancement of photosynthesis by adding ethanol (Fig. 4b) is very pronounced ((B* and C* > A). However, respiration started to increase on day 2 and exceeded photosynthesis (day 4). It can be assumed that the increased respiration in the presence of ethanol is accompanied by the intracellular formation of carbon dioxide. This is an additional substrate that stimulates the photosynthetic process. At the same time, it is precisely the production of glucose and reducing agents during photosynthesis that causes a cell culture to maintain a high respiration rate in the presence of ethanol and light (Fig. 4A), in contrast to darkness, when respiratory substrates are depleted (Fig. 2).

Autotrophic cells in darkness, deprived of light energy, are in a state of biological re-experiencing. For dark autotrophic cells of *G. sulphuraria*, an almost twofold smooth decrease in the rate of O_2_ uptake on the 4-th day of cultivation is evident due to depletion of respiratory substrates in the absence of photosynthesis (Fig. 5).

**Fig. 5.**
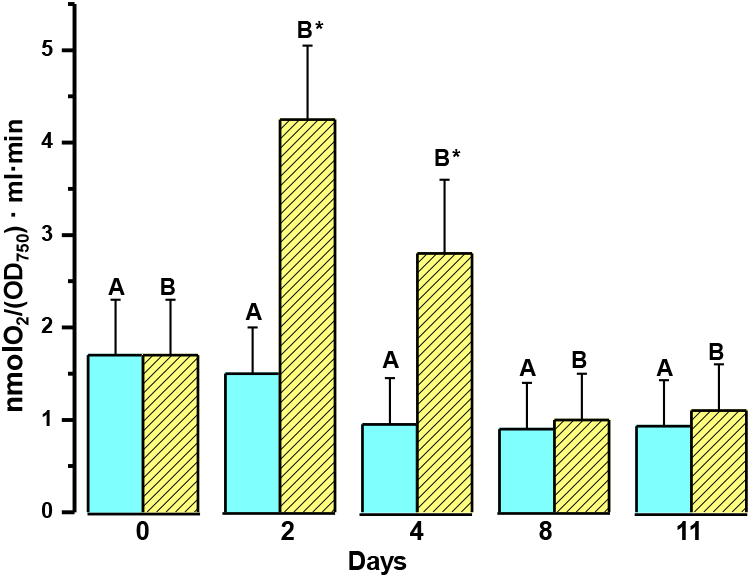
Oxygen uptake during dark cultivation of *G. sulphuraria*. Columns A correspond to dark surviving of autotrophic culture and columns B to the addition of 0.1% ethanol. Statistically significant differences are marked with *.

When ethanol is added to culture, respiration patterns change. In this case, oxygen uptake increased 2.5-fold on the second day of cultivation and then decreased just as rapidly to minimal values comparable to the control culture (Fig. 4). Since photosynthesis and growth of *G. sulphuraria* are absent in the dark (Fig. 2), as well as ethanol consumption (Fig. 3), it can be assumed that the short-term increase in respiration in this case is a cell response to mild ethanol stress, as ethanol enters the cells by diffusion.

### Absorption spectra *of G. sulphuraria*

The growth of the *G. sulphuraria* culture (see Fig. 2) was assessed by light scattering. However, light scattering is optically variable depending on the growth conditions and cell suspension density and significantly misrepresents the shape of the spectral curves. Therefore, the absorption spectra of cells exposed to light in autotrophic and ethanol-supplemented cultures were corrected over the entire visible spectral region from 400 to 750 nm by subtracting the light-scattering contribution (Fig. 6).

**Fig. 6.**
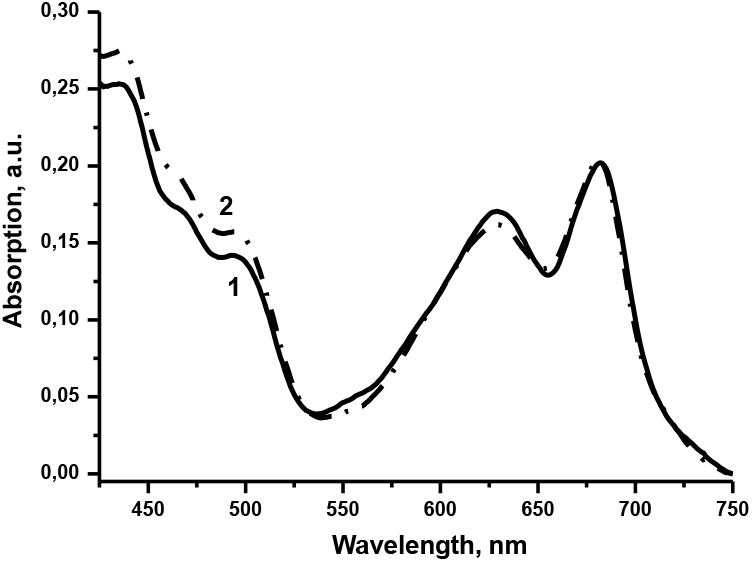
Absorption spectra of *G. sulphuraria* on the last day of cultivation in the light. Spectra are corrected for light scattering and normalised to the chlorophyll peak position at 678 nm. 1) photoautotrophic growth (solid line); 2) growth in the presence of 0.1% ethanol (dash dot line).

It can be seen that the overall shape of the spectra and the ratio of the intensities of the carotenoid (500 nm), phycobilisome (625 nm) and chlorophyll (678 nm) bands remain almost unchanged irrespective of the presence of ethanol. This can serve as an additional indicator of maintaining photosynthetic activity of *G. sulphuraria* when exposed to ethanol.

## Discussion

The combined data on the enhancement of cell culture growth in the presence of ethanol in the light, at the same time as its consumption from the growth medium, and the intensification of cellular respiration and photosynthesis, allows us to conclude that ethanol is metabolisable by the light-cultured *G. sulphuraria*.

In this paper, an ethyl alcohol concentration of 0.1% has been used to minimise the ethanol stress, as mentioned above. Due to methodological difficulties caused by varying stress-inducing concentrations, there are not too many microalgae for which ethanol assimilation has been described. Early reports are summarised in (Neilson and Lewin, 1974). Work in the last 15-20 years, as mentioned above, has been mainly concerned with a number of species of green algae and dinoflagellates. As mentioned above, research over the last 15-20 years has focused on a number of species of green algae and dinoflagellates. (See for example (de Swaaf et al., 2003; Ketsuda et al., 2006). To our knowledge, *G. sulphuraria* is the first red microalga to be added to the list. In the dark, there was no evidence of ethanol consumption from the culture medium due to its assimilation by *G. sulphuraria* cells (Figs. 2 and 3). Nevertheless, by diffusion, ethanol should enter the cells independently of illumination at a concentration equal to the culture medium. This is evidenced by an increase in cellular respiration in the dark (Fig. 5), which can be interpreted as a cellular response to the manifestation of the stress effect of ethanol. The rapid depletion of respiratory cellular resources not replenished by photosynthesis under dark conditions appears to explain the subsequent decline in respiratory activity (Fig. 5).

The inhibition of ethanol uptake by *G. sulphuraria* in the absence of light is an effect that has not been previously described for microalgae. However, it was observed that the unicellular green alga *Chlamydomonas mundana* is unable to assimilate acetate (a product of ethanol conversion) from the growth medium in the dark, but consumes it in the light (Eppley et al., 1963). Therefore, light dependence of exogenous ethanol consumption in *G. sulphuraria* culture is consistent with such findings. The existing cryptochrome photolyase system regulates the light-dependent activity of 35% of nuclear genes in the closely related *G. sulphuraria* species *Cyanidioschyzon merolae* (Asimgil and Kawakli, 2012; Tardu et al., 2016). In the case of *G. sulphuraria*, a similar photoreceptor mechanism appears to be associated with photosynthetic activity and stimulates ethanol assimilation. This is achieved, as in other microalgae, by the conversion of ethanol to acetaldehyde by ADH. This is followed by complete oxidation. The various forms of ADH are enzymes found in all three cellular kingdoms: archaea, bacteria and eukaryotes. Three non-homologous families of NAD(P^+^)-dependent ADHs are currently known, each with a large number of isozymes. Type I ADHs comprise Zn-dependent enzymes, type II is represented by short-chain ADH proteins, and type III corresponds to iron-containing ADHs (FeADHs). The three ADH families have arisen independently during evolution, have different protein structures and different molecular reaction mechanisms (Gaona-Lopez et al., 2016).

According to genomic data (Saura et al., 2022), *G. sulphuraria* contains glycerol dehydrogenase/iron-containing alcohol dehydrogenase (FeADH, Gasu_57960), a type III ADH. This evolutionarily early type of isozymes, is widely represented among prokaryotes and unicellular eukaryotic organisms (Gaona-Lopez et al. 2016). The data on ethanol metabolism by *G. sulphuraria* are fully consistent with the presence of this enzyme. This is also consistent with the presence of a gene for the second key enzyme in the ethanol conversion pathway, acetaldehyde dehydrogenase (Gasu_62010), which further converts the resulting acetaldehyde to acetate (Rossoni et al., 2019). By non-enzymatically acetylation the SH- and NH_2_-groups of proteins, it depresses mitochondrial respiratory chains and is an order of magnitude more toxic compound than ethanol itself (Bulloc, 1990). The cells are therefore ‘interested’ in the subsequent active metabolisation of acetaldehyde. The further pathway is well known: acetic acid is converted into acetyl-CoA, which enters the Krebs cycle with final CO_2_ release ((Saeki et al., 1997; Kornmann et al., 2003) and Fig. 1).

In addition to the main pathway - entry into the TCA cycle - acetyl-CoA can be used in lipid biosynthesis and in the glyoxalate cycle. In *G. sulphuraria*, the cytosolic highly branched floridean starch serves as a reserve polysaccharide (Stadnichuk et al., 2007; Martinez-Garcia et al., 2016), which is inconsistent with lipid synthesis at the expense of the incoming ethanol. The glyoxalate cycle is a modified TCA cycle that bypasses its decarboxylation step (Saeki et al., 1997). It is incompatible with the enhancement of photosynthesis in the presence of ethanol (Figure 6), which requires CO_2_. Both enzymes, ADH and ACDH, are considered to be active participants in cellular defence against oxygen stress and other stressors (Rossoni et al., 2019; Lin et al., 2021), and their role is not limited to ethanol oxidation alone. The light-induced enzymatic activity of ADH corresponds to the activation of photosynthesis in *G. sulphuraria* culture and consequently to the production of oxygen and the cell protection from oxygenating products.

The excess of photosynthesis when ethanol is added compared to the photoautotrophic culture probably has the same cause as known for *G. sulphuraria* grown on glucose. Photoheterotrophic growth during glucose addition is greater than the sum of chemoheterotrophic and photoautotrophic growth levels (Stadnichuk et al., 1998; Curien et al., 2021). This difference can be explained by the fact that during glucose oxidation, CO_2_ is transported from the mitochondria to the chloroplasts, where it is used as an additional carbon source, thereby enhancing photosynthesis.

Therefore, the increase in photosynthesis during ethanol assimilation can be explained by a similar utilisation of the formed CO_2_ as an end product of the oxidative process. In line with this speculation, the scheme (Fig. 1) can be extended to a scheme (Fig. 7) in which light activation of the key enzymes, ADH and ACDH, and the use of CO_2_, the end product of ethanol oxidation, as a substrate for photosynthesis play an essential role. Consuming ethanol in the light leads to a decrease in its intracellular concentration, thus reducing its stress effects and is simultaneously energetically beneficial. It can be assumed that a possible residual ethanol content in food waste should not limit, but rather stimulate the industrial use of *G. sulphuraria*.

**Fig. 7.**
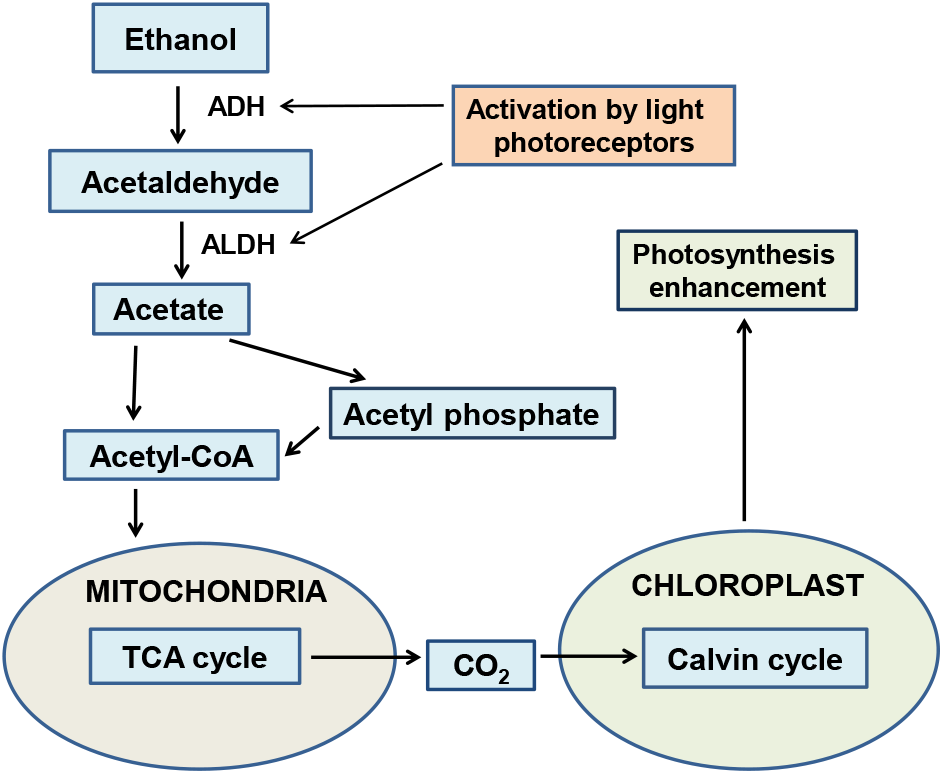
Extended scheme of ethanol oxidative assimilation assumed for *G. sulphuraria*, taking into account the light activation of ADH and ALDH and the utilisation of carbon dioxide formed in the TCA cycle in photosynthesis.

## Supplementary Information

The manuscript contains no supplementary material.

## Acknowledgments

The authors are indepted to Dr. V. Kevbrin for his assistance with ethanol chromatography determination.

## Data availability

The datasets generated during the current study are available from the corresponding author on reasonable request.

## Author contribution Declarations

I.S. and Y.B. contributed equally to the experimental work, analyzing the data and writing the manuscript.

